# Single-cell transcriptomics reveals a differential response of human bronchial epithelial cell-types to cadmium chloride

**DOI:** 10.64898/2026.02.23.707356

**Authors:** Fadi Abou Choucha, Rafaël Lopes Goncalves, Thomas Hermet, Jessica Mille, Laetitia Guardini, Mariem Benkhedher, Caroline Lacoux, Marine Gautier-Isola, Baharia Mograbi, Jérémie Roux, Françoise Cottrez, Bernard Mari, Hervé Groux, Claude Pasquier, Roger Rezzonico, Georges Vassaux

## Abstract

Exposure of cells or tissues to chemical compounds can be analyzed through transcriptomic signatures, which can be used to classify chemical agents. This information can also enrich Adverse Outcome Pathways (AOP). Transcriptional signatures have generally been obtained using “bulk” analysis, by which the global gene expression pattern of an entire tissue is determined. Although this approach has been useful in toxicology, some information is lost, especially when tissues containing multiple cell types are considered. With the advent of single-cell transcriptomics (scRNA-seq), it is now possible to obtain higher resolution, cell type–specific responses in complex tissues. The aim of the present study was to evaluate the added value of scRNA-seq in analysis of the acute response of human bronchial epithelial cells grown at the air/liquid interface (ALI) to a known toxic compound, CdCl_2,_ with well described transcriptional signatures of exposure. Fully differentiated mucocilliary epithelia obtained from three independent donors were exposed to 10 µM CdCl_2_ and scRNA-seq analysis was performed on a total of 18255 cells to obtain cell type–specific signatures. Our results show that the contribution of each cell type to the overall transcriptomic bulk response varies. For example, the classical heavy metal detoxification response was only detected in multiciliated and secreting cells, while absent in basal cells. The data demonstrate that scRNA-seq provides high-resolution transcriptional signatures with unexpected features. This added information is likely to have implications for the refinement of AOPs and could serve as a basis for a new generation of tests in predictive toxicology.

## INTRODUCTION

The US National Research Council defines toxicogenomics as the combination of “toxicology with information dense genomic technologies to integrate toxicant-specific alterations in gene, protein and metabolite expression patterns with phenotypic responses of cells, tissues and organisms”. Within this framework, transcriptomics has provided invaluable information on the interaction and mechanisms of action of compounds in a wide variety of biological systems, ranging from whole organisms, organs tissues and cell cultures. For example, in the field of carcinogenicity, transcriptomics has led to the establishment of fingerprints, transcritptional signatures, that provide a rationale to classify compounds and predict their mode of action (reviewed in (David 2020). These transcriptional signatures allowed for the successful classification of genotoxic and non-genotoxic carcinogens in a blinded study (Hamadeh et al. 2002) and the characterization of hepatic carcinogens showing ambiguous genotoxicity results (Kossler et al. 2015).

The methods used to generate transcriptional signatures have evolved from fluorescence detection methods such as microarrays to next generation sequencing, which counts reverse transcribed cDNAs (Alexander-Dann et al. 2018; David 2020). The platforms used to exploit the raw data are now well established and tables of genes differentially expressed in response to exposures of cells or tissues to chemical compounds have been compiled in multiple publicly accessible databases (Alexander-Dann et al. 2018). The information accumulated provides a unique set of resources to characterize the mechanism of action of a potential toxicant and can also be used to enrich adverse outcome pathways (AOPs). In addition, once a signature of a specific type of toxicity has been established, qRT-PCR–based tests can be set up and performed routinely at relatively low cost, to monitor the expression of a relevant set of genes in the response of a tissue to chemical compound. Such PCR-based assays can provide a rationale for classifying potential toxicants/irritants, as in the case of the SENS-IS test assessing skin sensitization (Cottrez et al. 2016; Cottrez et al. 2025).

Overall, transcriptional signatures have been obtained using “bulk” RNA preparations, in which a whole sample is homogenized and total RNA is prepared. Although this approach may provide sufficient resolution when studying the effects of compounds on a single cell type with limited heterogeneity, some information is lost when tests are conducted on complex tissues or cultures composed of multiple cell types. This limitation is particularly important in the context of broad international efforts to develop of highly sophisticated, multi-cell-type in vitro culture models to reduce or replace animal experimentation in drug testing (Stresser et al. 2023).

The development of single-cell transcriptomics (scRNA-seq) has triggered international initiatives leading to the generation of cell atlases of all the tissues in the human body (https://www.humancellatlas.org/). These initiatives have now been extended to the generation of atlases of diseased tissues see, for example, (Sikkema et al. 2023) for the Human Lung Cell Atlas. ScRNA-seq has also been used to describe changes associated with treatment tolerance and response to therapeutic agents in cancer (Aissa et al. 2021; Cambien et al. 2020; Ho et al. 2018), and virus-host interactions (Ratnasiri et al. 2023). The use of single-cell transcriptomics in toxicology and/or environmental pollution has been advocated in multiple reviews and commentaries (David 2020; Filipovic et al. 2024; Zhang et al. 2019) but characterization of human-cell-type heterogeneity in response to toxicants, and formal demonstration that scRNA-seq can actually provide unique information useful in the context of toxicology or AOPs, are relatively scarce (Chu et al. 2024; Filipovic et al. 2024; Gupta et al. 2022; Khan et al. 2023; Matchett et al. 2024).

Human organotypic airway epithelia derived from primary bronchial basal cells cultured at an air-liquid interface (ALI) represent a well-documented *in vitro* tissue model for physiological, pharmacological, and toxicological studies in lungs (Baxter et al. 2015; Cao et al. 2021). These cultures are the gold standard in human lung studies and represent a useful tool for reducing or replacing animal experimentation. Human ALI tissues have been widely used to study the effects of pollutants, chemicals, and nanoparticles (Petpiroon et al. 2023). These cultures recapitulate the cell populations found in native airway tissues, namely basal, secretory (club and goblet cells), and multiciliated cells (Ruiz Garcia et al. 2019). In addition, markers for these cell types suitable for droplet-based scRNA-seq have been documented (Ruiz Garcia et al. 2019).

Tobacco smoke is a major source of cadmium (Cd) exposure in the human populations (https://www.atsdr.cdc.gov/ToxProfiles/tp5-p.pdf). Cadmium is classified as a Group I carcinogen in humans and a major health concern by the World Health Organization. The chemical properties of Cd^2+^ are close to those of calcium (Ca) and zinc (Zn). As a result, Cd^2+^ interferes with cellular signalling events that implicate Ca^2+^ and with the function of protein that use Zn^2+^ as a co-factor (Forcella et al. 2022). At the cellular level, Cd^2+^ has been shown to affect various mechanism that include DNA repair, activation of tumor suppressors such as p53 (Chen et al. 2019; Lee and Thévenod 2020; Oldani et al. 2020; Urani et al. 2015). In term of bulk transcriptional responses, the Gene Ontology (GO) terms obtained upon analysis of the differentially expressed genes in Cd^2+^-exposed cell lines relate to metal ions responses and detoxification processes. This GO signature is observed in human cells of different anatomical origins, including the human lung adenocarcinoma cell line A549 (Forcella et al. 2022; Gasser et al. 2022a; Jomova et al. 2025).

In the present study, intended as a paradigm to illustrate the potential added value of scRNA-seq in toxicology, we used scRNA-seq to investigate the effects of acute CdCl_2_ exposure on different cell types present in human organotypic airway epithelial ALI cultures.

## MATERIALS AND METHODS

### Cell Culture

In vitro tissue model of human bronchial airway epithelium (MucilAir™-Bronchial) cultured at the ALI (MB-ALI) were purchased from Epithelix (https://www.epithelix.com/). Epithelix provides ready-to-use tissues reconstituted with primary bronchial cells that have been obtained according to the legal and ethical requirements of the country of collection, i.e. with the approval of an ethics committee and with anonymous written consent from the donor or nearest relative. These 3D cultures were obtained from three independent nonsmoking donors: Donor1: male, Caucasian, 35 years old; Donor 2: male, Hispanic, 42 years old; Donor 3: female, African American, 61 years old. Upon reception of ALI cultures, the medium (Mucilair culture medium) was changed and cells were left to recover for one week in an incubator (5% CO_2_ at 37°C) before experimentation. The BEAS-2B cell line (ATCC CRL-3588) was isolated and immortalized from normal human bronchial epithelium and routinely cultured in Dulbecco’s modified Eagle medium (DMEM) supplemented with 10% heat inactivated fetal bovine serum (Pan Biotech).

### CdCl_2_ exposure of cells and toxicity assays

MB-ALI were incubated with Mucilair medium alone or containing DMSO (0.1%) or CdCl_2_ (10µM) in 0.1% DMSO (700 µL on the basal side and 200 µL on the apical side). BEAS-2B cells were incubated in complete medium containing DMSO (0.1%) or various concentration of CdCl_2_ in 0.1% DMSO. Twenty-four hours after adding CdCl_2_ on MB-ALI or BEAS-2B cultures, 200 µL of medium was collected and Lactate dehydrogenase (LDH) release was measured using the CyQUANT LDH cytotoxicity assay (Invitrogen, as an indicator of cell death.

### ScRNA-seq

After six hours of exposure, the MB-ALI inserts were washed twice with 1 x HBSS (Gibco) (700 µL at the basal side and 200 µL at the apical side). Tissues were then digested overnight at 4°C with a solution of 0.15% pronase (Sigma Aldrich/Merck) in 1 x HBSS. The next day, cells were dissociated mechanically and the reaction was stopped by adding 1 mL of a 2% bovine serum albumin (BSA) solution. The cell suspension was then filtered on a strainer (40 µm), and the monocellular cell suspension was washed twice in a 1% BSA solution. Fixation and permeabilization of the cells were performed according to the protocol of the Evercode cell fixation kit (PARSE bioscience). Cells were then stored at – 80°C. After thawing, cells were counted. Barcoding and generation of the libraries were performed as described in the Evercode WT v2 kit (Parse bioscience). The scRNA-seq libraries were sequenced as indicated in the Evercode WT v2 kit (Parse bioscience) manual instruction.

### Generating single-cell gene expression matrices, quality control and filtering

Raw reads were processed to generate gene expression matrix using the split-pipe pipeline (v1.0.3p) from Parse Biosciences (https://support.parsebiosciences.com/hc/en-us/categories/360004765711-Computational-Support). The GRCh38/hg38 human reference genome was used to map and quantify the gene expression. The human genome reference files were downloaded from the Ensembl database (https://ftp.ensembl.org/pub/release108/fasta/homo_sapiens/dna/Homo_sapiens.GRCh38.dna.primary_assembly.fa.gz; https://ftp.ensembl.org/pub/release-108/gtf/homo_sapiens/Homo_sapiens.GRCh38.108.gtf.gz,). The following parameters: --mode all; -- chemistry v2; --kit WT were used on the split-pipe tool (v1.0.3p).

The expression matrix was loaded into the R package (v.4.2.0) using Seurat (4.4.0) (https://satijalab.org/seurat/articles/install_v5.html) for quality control and downstream analyses. Cells with greater than 5 median absolute deviations (MAD) and cells with >20% of their transcriptome of mitochondrial origin were removed. Gene counts values were then normalized and transformed with NormalizeData Seurat function, which normalizes the total counts for each cell by the total counts, multiplies this by a scale factor of 10,000, then log-transformed the result. Principal component analysis (PCA) as well as UMAP using the first 30 principal components were performed. UMAPs of individual samples were inspected before integration.

### Data integration

Individual samples were integrated in Seurat using the reciprocal PCA (RPCA) pipeline to remove batch effects. The ‘SelectIntegrationFeatures’ function was applied to choose the 3000 features ranked by the number of datasets they were detected in. Next, the ‘FindIntegrationAnchors’ function selected a set of anchors between different samples using the top 30 dimensions from the RPCA to specify the neighbour search space. The three control samples (Ctrl_DMSO_1pc_D1, Ctrl_DMSO_1pc_D2, Ctrl_DMSO_1pc_D3) were specified as a reference. ‘IntegrateData’ was then applied to integrate the samples using the pre-computed anchors and the integrated dataset was scaled using ‘ScaleData’. PCA and UMAP dimension reduction based on the top 30 principal components were performed. Nearest-neighbour graphs using the top 30 dimensions of the PCA reduction were calculated and clustering was applied with a resolution of 0.2 and 0.4.

### Cell-type identification and differential analysis

Differential gene expression (DGE) in each cluster was computed with ‘FindAllMarkers’ on ‘RNA’ assay. The cell types were identified by manually curating the cluster DGE with the reported literature, such as the single-cell lung atlas (DOI: 10.1165/rcmb.2018-0416TR) and Human Cell Atlas (https://www.humancellatlas.org). We compared our cluster DGE with the Azimuth (https://azimuth.hubmapconsortium.org) and Celltypist (https://www.celltypist.org) shared high expression in scRNA-seq data. The final results were manually examined to ensure the correctness of the results. The three major cell types were chosen by initial exploratory inspection of the differentially expressed genes (DEGs) of each cluster combined with literature study. For visualization purposes, expression scores were plotted in UMAP embeddings or violin plots using log-normalized values. Unless otherwise noted, we calculated the proportions of cell types per sample, and compared the medians of the two groups to identify proportion differences.

### Pseudobulk and gene set enrichment analysis

To test the variation across samples, we created the H5AD file from annotated data with three cell types. Pseudobulk profile for each donor, celltype and condition, was obtained by using the decoupler (v3.8, https://github.com/HBNetwork/python-decouple) function ‘dc.get_pseudobulk’ on normalized count data. Next, DEG was computed by comparing the gene expression of each cell type from each condition against control (DMSO) for the three donors. The framework of DESeq2 implemented in decoupler tool was used for this experimental design. Differentially-expressed genes were used to perform enrichment analysis using ingenuity pathway analysis (IPA) tool (QIAGEN Inc., https://digitalinsights.qiagen.com/IPA) and Reactome Pathway Database gene sets (Griss et al. 2020).

### Gene Regulatory Network Inference Using SCENIC

We reconstructed gene regulatory networks (GRNs) and inferred transcription factor (TF) activity at the single-cell level using pySCENIC (*scenic.aertslab.org*), a Python-based implementation of the SCENIC method (Aibar et al. 2017). Single-cell RNA sequencing (scRNA-seq) data for CdCl_2_ and DMSO controls were analyzed using the *Homo sapiens* (hg38) RefSeq r80 reference genome. Gene co-expression modules were identified with GRNBoost2 within pySCENIC, which employs gradient boosting machines to detect regulatory interactions based on expression patterns, generating an initial gene-gene network. Cis-regulatory motif enrichment was performed for each co-expression module using SCENIC’s motif databases (*resources.aertslab.org/cistarget/*): Distal Elements ±10 kb from the transcription start site (TSS), and Proximal Promoters 500 bp upstream to 100 bp downstream of the TSS. RcisTarget assessed motif overrepresentation in the upstream regions of module genes. Regulons were defined by integrating co-expression and motif enrichment data, retaining only those with ≥10 genes and a Normalized Enrichment Score (NES) ≥3.0, and TFs with direct binding site annotations. We quantified regulon activity per cell using AUCell (Aibar et al. 2017) by calculating the Area Under the Curve (AUC) for each regulon. The AUC scores were normalized, scaled to 0–1, and binarized to classify cells as “active” or “inactive” for each regulon.

### Differential Regulon Activity Analysis

Using the Scanpy toolkit (*scanpy.readthedocs.io*) (Wolf et al. 2018), we normalized the AUC matrix by the maximum AUC per regulon and scaled activities between 0 and 1. Cells were grouped by cell type and treatment, using control conditions as references. Differential regulon activity was assessed with the Wilcoxon rank-sum test, and p-values were adjusted using the Benjamini-Hochberg method. All regulons were included to minimize bias, with significant differential activity defined as adjusted p-value < 0.05 and log fold change > 0.5.

### Gene Set Enrichment and Network Analysis

To analyze functional relationships among genes in identified regulons, we performed gene set enrichment analysis using gProfiler (Raudvere et al. 2019) and visualized the results in Cytoscape 3.10 (cytoscape.org). We selected regulons showing differential activity (p-value < 0.05) in at least one cell type and extracted their target genes for each cell type. Enrichment analysis was conducted against GO Biological Process and Reactome databases, applying Benjamini-Hochberg FDR correction (FDR < 0.05) and filtering for gene sets containing 10-500 genes. The top 50 enriched pathways were exported as a GMT file and imported into Cytoscape’s Enrichment Map app. The final network was filtered to display only gene sets with FDR q-value < 0.01, and unconnected or minimally connected nodes (≤1 edge) were hidden to enhance network clarity.

### Single-molecule RNA Fluorescence in Situ Hybridization (RNA FISH)

RNA FISH on formalin-fixed paraffin embedded (FFPE) sections were performed using the RNAscope Multiplex Fluorescent assays V2 (Advanced Cell Diagnostics, Hayward, CA) according to the manufacturer’s protocol. IL1RN-C1, SYTL5-C1, WIPF3-C1, MT1G-C1, commercial probes were used to detect these RNAs. Briefly, MB-ALI cultures were formalin-fixed and embedded in paraffin. FFPE tissue sections (7 µm) were deparaffinized, dehydrated, treated to inhibit peroxidase activity, submitted to target retrieval treatment and digested with protease Plus reagent for 30 min at 40°C before probes hybridization. The signal was revealed by sequential incubations with amplifier reagents and dyes according to manufacturer’s protocol. Co-immunolabelling with mouse anti-Acetyl-alpha tubulin (Sigma, 6-11B-1) and Alexa Fluor® 488 anti-cytokeratin 5 (Abcam, EP1601Y), was performed following RNA-FISH experiments. Briefly, sections were blocked for 1h in Animal-Free blocking solution (Cell signalling) and incubated overnight at 4°C with primary antibodies. After washing with PBS, anti-mouse Alexa Fluor 647 (ThermoFisher Scientific) was added for 1h at room temperature. DAPI was used to counterstain nuclei and coverslips were mounted onto glass slides using either Fluoromount-G (ThermoFisher Scientific). Fluorescence was visualized and captured on a Zeiss LSM780 confocal microscope.

## RESULTS

We first evaluated the toxicity of CdCl_2_ on BEAS-2B pulmonary epithelial cells. **Suppl. Fig.1A** indicates that a dose-dependent toxicity is observed, with a concentration of 10 µM inducing around 5% of cell death on these cells. This concentration was tested on MB-ALI cultures and **Suppl Fig.1B** shows that 10 µM of CdCl2 led to 1% of cell death after 24 hours of exposure This concentration was chosen in the rest of the study to make sure that cadmium signatures after six hours of exposure were at their maximum, without apoptotic interferences.

To characterize the response of the different epithelial cell-types in the MB-ALI culture from three individual donors, 10 µM of CdCl_2_ was added on both the apical and basal sides. After six hours of treatment, monocellular suspensions were generated. The cells were then fixed, permeabilized and subjected to the PARSE Bioscience barcoding and library construction protocols, followed by sequencing (**Figure 1A**). A standard analysis was performed using cells with a percentage of mitochondrial genes below 20 %. On the uniform manifold approximation and projection (UMAP) plot generated, the cells clustered into three main clusters: basal, secretory and multiciliated cells (**Figure 1B**), based on the expression of cell-type-specific markers (**Figure 1C**). The control (DMSO)- or CdCl_2_-treated cells were distributed evenly across the different cell-types (**Figure 1D**) and the proportion of cells in the two conditions was comparable (**Figure 1E**). In a first instance, a “pseudobulk as bulk” analysis was performed, in which all the cell-types were pooled. This type of analysis is equivalent to a bulk RNA sequencing. The differential expression analysis of cells treated with CdCl_2_ versus solvent (DMSO) provided a transcriptomic signature (**Figure 1F**). This signature is composed of 242 genes up- or down-regulated. In particular, genes involved in metal ion homeostatis and detoxification were upregulated. The complete list of modulated genes is presented in **Suppl. Table 1**. This transcriptional signature is homogeneous across the three donors, as shown in the heatmap in **Figure 1G**, representing the top 50 modulated genes. Ingenuity Pathways analysis™ (IPA) of the canonical pathways modulated by CdCl_2_ exposure demonstrates a classic response to heavy metals (term IPA metallothioneins bind metals), a modification of the extracellular matrix (ECM, terms syndecan interactions, extracellular matrix organization, and integrin cell surface interactions), and a cellular stress/anti-oxidant response (term KEAP1-NFE2L2 pathway) (**Figure 1H**). IPA also provided a list of upstream regulators modulated upon CdCl_2_ exposure. They include a positive regulation of inflammation (term TNF), of tissue repair (terms PDGF-BB, FGF2, EGF, TGFb1), of exposure to heavy metals (term metoxalin gadolinium), and of the presence of reactive oxygen species (ROS) (term hydrogen peroxide) (**Figure 1I**). Altogether, these responses are consistent with the expected transcriptional signature linked to exposure of epithelia to CdCl_2_ (Forcella et al. 2022; Gasser et al. 2022a; Gasser et al. 2022b).

**Figure 1.**
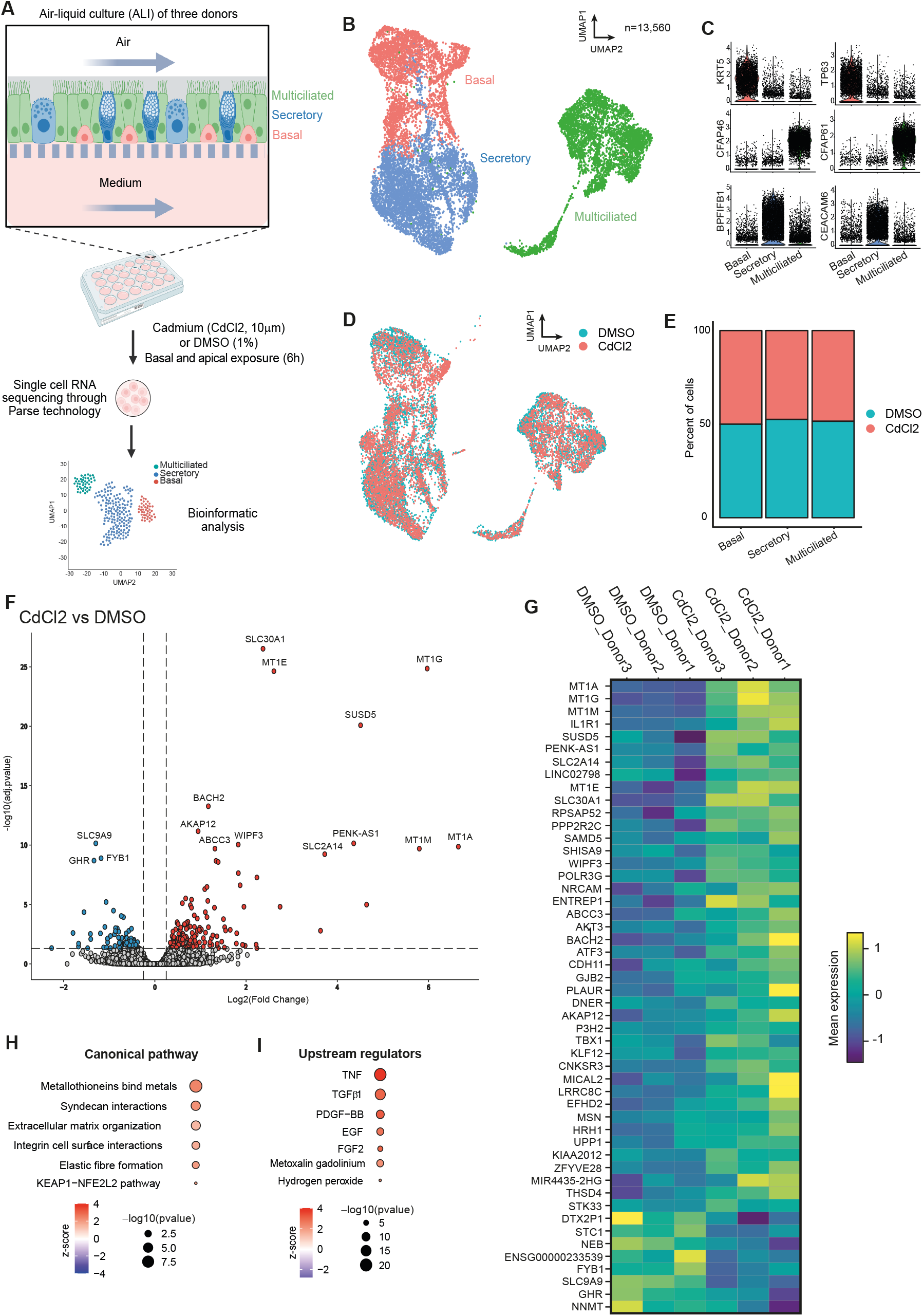
Experimental design and “pseudobulk as bulk” transcriptional analysis of cadmium exposure in human bronchial epithelial cells. (A) Experimental design. (B) Uniform manifold approximation and projection (UMAP) plot presenting the different cell-types in the dataset. (C) Violin plots presenting the expression of cell-type-specific markers for basal, secretory, and multiciliated cells. (D) UMAP presenting the distribution of cell types in the different conditions. (E) Proportions of the different cell-types in control or CdCl_2_-exposed cultures. (F) Volcano plot of differentially expressed genes (CdCl_2_ versus control) in the “pseudobulk as bulk” analysis. (G) Top 50 modulated genes (CdCl_2_ versus control) across the three donors. (H) Ingenuity Pathway Analysis (IPA) of the canonical pathways and upstream regulators (H) modulated by CdCl_2_ exposure.

We next examined the effects of CdCl_2_ exposure on the different cell-types in the MB-ALI cultures. **Figure 2A** presents the number of cells in the different cell-types. Basal, secretory and multiciliated cells represent 27.3%, 40.5% and 32.2% of the cells, respectively. These proportions can be compared to the proportion of differentially-expressed genes in each cell types: 65%, 15.3% and 19.7%, for basal, secretory and multiciliated cells, respectively (**Figure 2B**). This observation suggests that basal cells are the most responsive to CdCl_2_ exposure. Differential expression analyses of cells treated with CdCl_2_ versus solvent (DMSO) performed on basal, secretory and multiciliated cells are presented in the vulcanoplots in **Figures 2C, 2D and 2E** (the complete list of modulated genes in basal, secretory and multiciliated cells is presented in **Suppl. Table 2, 3 and 4**, respectively). As for the “pseudobulk as bulk” (**Figure 1F**), these transcriptional signatures are homogeneous across the three donors, as shown in the heatmap in **Figures 2C, D and E**, representing the top modulated genes. Canonical pathways and upstream regulators affected were evaluated using IPA. Analysis of modulated canonical pathways showed that the classic response to heavy metals is restricted to secretory and multiciliated cells but absent from basal cells. By contrast, cytoskeletal reorganization (IPA term Rho GTPase cycle) is activated only in these basal cells. Finally, a cellular stress response and a hypoxic signature are detected only in multiciliated cells (term NFE2L2 regulating antioxidant/detoxification enzymes and HIF1a signalling, respectively) **(Figure 2F)**. In term of upstream regulators, a strong, positive regulation of inflammation (term TNF) was observed in all three cell types. Tissue repair (terms PDGF-BB, FGF2, EGF, TGFb1) was essentially detectable in basal and secretory cells. Exposure to exposure to heavy metals (term metoxalin gadolinium) was strongly activated in secretory and multiciliated cells. Finally, the presence of ROS (term hydrogen peroxide) in basal and multiciliated cells (**Figure 2G**).

**Figure 2.**
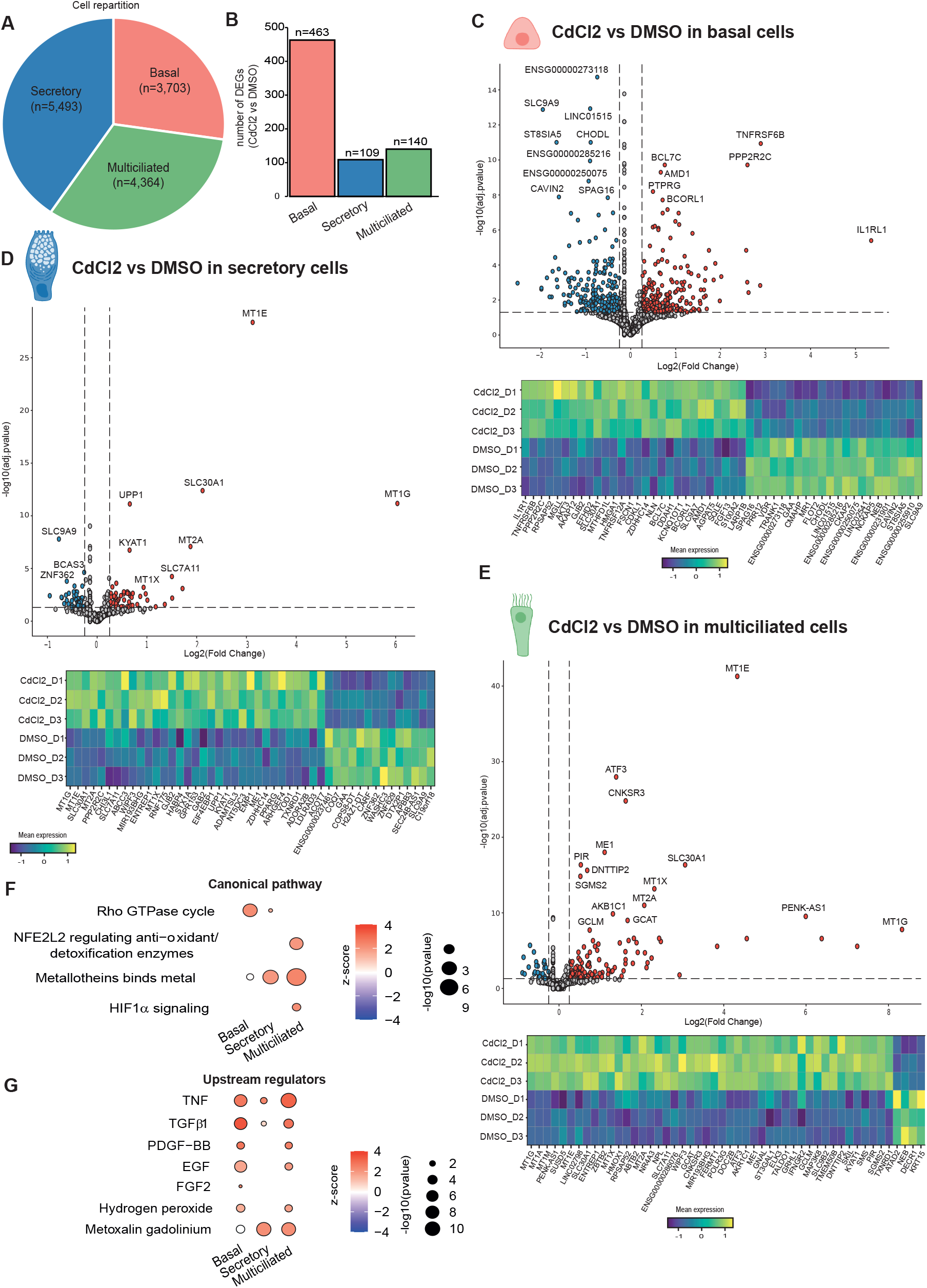
Cell-type specific transcriptional analysis of cadmium exposure in human bronchial epithelial cells. (A) Distribution of the different cell-types in the dataset. (B) Number of genes differentially modulated CdCl_2_ versus control) in the different cell-types. Volcano plots of and heatmap per donor of the differentially-expressed genes (CdCl_2_ versus control) in the basal (C), secretory (D) and multiciliated cells (E). Ingenuity Pathway Analysis (IPA) of the canonical pathways (F) and upstream regulators (G) modulated by CdCl_2_ exposure in the different cell-types.

In this study CdCl_2_ was dissolved in DMSO and the final concentration of DMSO on the cells was 0.1%. As this solvent is not neutral, we analysed the transcriptomic signature of 0.1% DMSO on MB-ALI cultures (**Suppl. Fig 2**). As for the CdCl_2_ versus 0.1% DMSO analysis (**Figure 1**), the UMAP plot presents three main clusters: basal, secretory and multiciliated cells (**Suppl. Fig 2A**). The cells were distributed evenly across the different cell-types (**Suppl Figure 2B**) and the proportion of cells in the two conditions was comparable (**Suppl. Fig 2C**). Pseudobulk as bulk as well as cell-type specific responses are presented in **Suppl Fig 2D-F** and the differentially-expressed genes are listed in **Suppl. Tables 5-8**. Of note, and unlike in the CdCl_2_ versus solvent analysis (**Figures 1 and 2**), heterogeneity was observed across donors (**Suppl Fig 2D-F**), suggesting a weak response of the cells to 0.1% DMSO. When the number of genes modulated is concerned, basal cells were the least responsive. Ingenuity Pathways analysis™ showed canonical pathway were essentially inhibited, in particular the RHO GTPase cycle (in the bulk as well as secretory and multiciliated cells, **Suppl. Fig 2H**), as well as the NRF2-mediated oxidative stress response (multiciliated cells, **Suppl. Fig 2I**). In term of upstream regulators, the TNF, TGFb FGF2, EGF and hydrogen peroxide were inhibited in secretory and multiciliated cells (**Suppl. Fig 2I**). Considering that these pathways and upstream regulators were induced in the CdCl_2_ versus 0.1% DMSO analysis (**Figures 1 and 2**), the utilization of DMSO as a solvent is likely to have blunted these responses. In addition, an anti-inflammatory response was also detected in bulk and all cell-types (upstream regulator LPS, **Suppl. Fig 2I**). The DMSO signatures obtained are consistent with the known effects of DMSO on cellular systems (Dubois-Pot-Schneider et al. 2022; Sanmartín-Suárez et al. 2011; Santos et al. 2003). Finally, we compared the genes differentially-expressed in DMSO versus medium and CdCl_2_ versus DMSO in the bulk and the three cell types. The Venn diagrams in **Suppl. Fig.2J** shows that very few genes were commonly modulated in the two analysis. Altogether, although the use of 0.1% DMSO as solvent in not neutral and has blunted the oxidative stress responses and the cytoskeletal reorganization (IPA term Rho GTPase cycle), it has not affected the differential activation of the classical heavy metal response in secretory and multiciliated cells compared to basal cells.

These data (**Figures 1 and 2**) were obtained using the pseudobulk method of analysis. Pseudobulk is a computational approach that mimics the structure of bulk RNA-seq data by averaging counts from individual cells in a given population. To verify the conclusions of the pseudobulk analysis, we performed an analysis based on the single-cell network interence and clustering (SCENIC) methodology, a computational method that infers gene regulatory networks and identifies cell states based on regulon activity (Aibar et al. 2017). In a first instance, the whole dataset was analysed as a bulk, without accounting for the different cell types present in the MB-ALI culture. **Figure 3A** shows that the 11 top regulons differentially-modulated upon CdCl_2_ exposure (absolute value of activity LogFC > 0.5). This pattern of regulon modulation is consistent with a state of epithelial activation and plasticity, such as that seen in wound healing, regeneration, or partial EMT. The known or plausible role of these 11 top-regulons modulated are presented in **Table 1**. The activity of these 11 top regulons was analysed in each cell types of the MB-ALI culture. **Figure 3B** shows that, globally, these regulons are modulated in a similar way in the three cell-types and in a way that matches regulon activation observed in bulk analysis (**Figure 3A**). One notable exception is the regulon MAFG, which is hardly affected in basal cells and highly activated in secretory and multiciliated cells (**Figure 3B**).

**Table 1:** Role of the top-regulons modulated by CdCl_2_ exposure. The known or plausible role of the 11 top-regulons modulated in epithelial plasticity and epithelial/mesenchymal transition (EMT) are presented. This table was generated using Chat GPT, with the question, “what would be the physiological and regulatory roles of the selected transcription factor/regulons in epithelial cells, focusing on states associated with EMT, stress adaptation, and regeneration”.

| Regulon / Transcription Factor | Known or plausible role in epithelial / EMT / plasticity | Comment / Supporting Evidence / Mechanistic notes |
| --- | --- | --- |
| ZEB1 | Master EMT inducer — represses epithelial genes (E-cadherin/CDH1, tight junctions), activates mesenchymal and stemness programs. | Drives epithelial-to-mesenchymal transition (EMT), invasion, and loss of polarity. Canonical EMT factor. |
| BCL3 | Modulates NF-κB signaling; influences epithelial proliferation and inflammatory responses. | In intestinal and mammary epithelium, BCL3 can restrain excessive NF-κB-driven inflammation and modulate Cyclin D1; dysregulation can promote tumorigenesis. |
| HMGA1 | Chromatin architectural factor; promotes chromatin plasticity, proliferation, and dedifferentiation. | Upregulated in proliferative or stem-like epithelial states; supports chromatin reorganization during stress or transformation. |
| MAFG (with NRF2) | Contributes to cellular defense against heavy metal toxicity by regulating the expression of antioxidant and phase II detoxification enzymes. | NRF2–MAFG heterodimers bind Antioxidant Response Elements; Activation mitigates oxidative stress and metal-induced cellular damage. |
| FOSL1 (FRA-1) | Part of AP-1 complex (FOS family); mediates proliferation, motility, invasion, and EMT. | Induced by MAPK/ERK signaling; promotes migration and extracellular-matrix remodeling. Elevated in wound repair and carcinoma progression. |
| CREB5 | cAMP/PKA-responsive TF; regulates stress, metabolism, and survival pathways. | May integrate metabolic and stress signaling in epithelial repair contexts; can cooperate with other bZIP TFs (e.g., ATF family). |
| DLX3 | Homeobox factor regulating differentiation, particularly in epidermal and hair follicle lineages. | Controls keratinocyte differentiation and barrier formation; its activation suggests morphogenetic remodeling or terminal differentiation pathways. |

**Figure 3.**
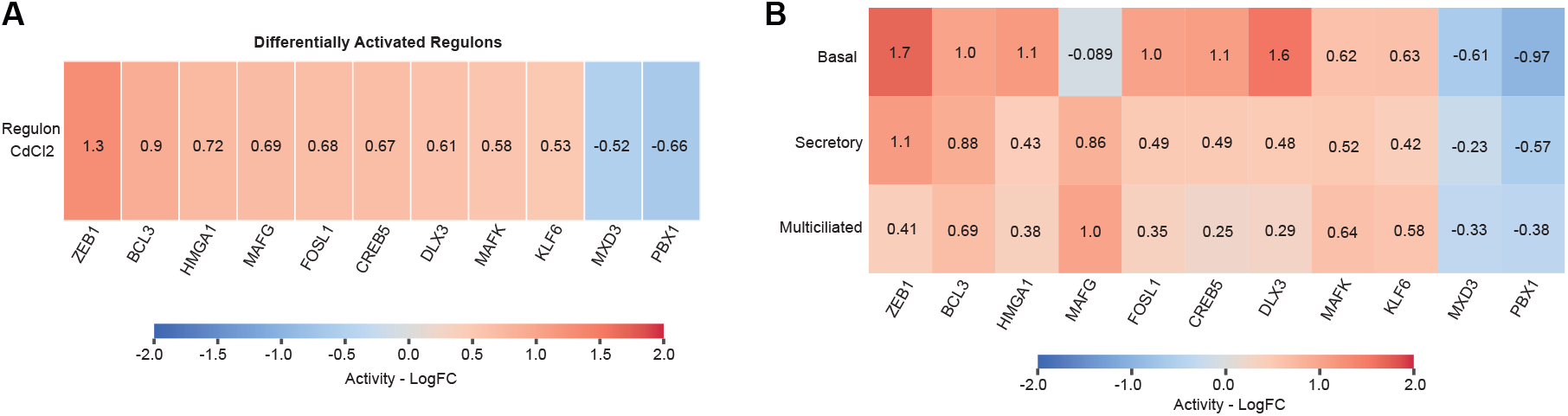
Regulon Activity and Expression in Response to CdCl_2_ Treatment. (A) Differentially Activated Regulons in CdCl_2_-Treated Cells. Heatmap displays regulons with significantly increased activity (logFC > 0.5; p-value < 0.05) in CdCl_2_-exposed cells compared to DMSO control. Regulon activity is quantified as log fold change (logFC) of activity scores computed using the AUCell algorithm (see Methods). (B) Cell Type-Specific Regulon Activity. Heatmap shows differential regulon activity (logFC > 0.5 in at least one cell type; p-value < 0.05) across basal, multiciliated, and secretory cell populations. Red indicates increased activity (positive logFC), while blue indicates decreased activity (negative logFC). (C) MAFG target gene expression across cell types. Dot plot shows expression of MAFG transcription factor target genes in basal, secretory, and multiciliated cells. Dot size represents the percentage of cells expressing each gene within a given cell type. Color intensity (light to dark red) indicates average normalized expression level, with darker shades representing higher expression. (D) Ingenuity Pathway Analysis (IPA) of the canonical pathways associated with the subset of genes composing the MAFG regulon and modulated by cadmium exposure.

Activation of the MAFG regulon is linked to responses to metal ions and stresses (**Figure 4A**). In the context of this study, MAFG, as a dimerization partner of NRF2, contributes to cellular defence against heavy metal toxicity by regulating the expression of antioxidant and phase II detoxification enzymes (Hirotsu et al. 2012; Katsuoka and Yamamoto 2016). The fact that this regulon is selectively activated in secretory and multiciliated cells is consistent with the conclusions of the pseudobulk analysis (**Figure 2**). To elucidate cell type-specific transcriptional responses to cadmium exposure, we performed enrichment map analysis on highly active regulons (activity logFC > 0.5) from basal, secretory, and multiciliated airway epithelial cells. The resulting network revealed both shared (multicolored nodes) and cell type-specific (single-color nodes) pathway activation patterns (**Figure 4B and Supplementary Figure 3**). **Figure 4B** illustrates the cadmium-response subnetwork, with three functionally distinct clusters. The upper cluster, predominantly driven by secretory (blue) and multiciliated (green) cells, was enriched for detoxification pathways including cellular response to cadmium ion, response to toxic substances, detoxification of inorganic compounds, and response to metal ions. The dense interconnectivity among these pathways suggests coordinated cellular defence mechanisms against cadmium. Both cell types also shared activation of the cellular response to oxygen-containing compounds pathway, indicating convergent stress responses in differentiated airway epithelia. In contrast, the lower subnetwork region revealed cell type-specific adaptations. Basal cells (red) uniquely activated transmembrane receptor protein tyrosine kinase signalling and growth factor response pathways, suggesting prioritization of proliferative and regenerative programs. The full network is presented in **Supplemental Figure 3**. These findings demonstrate that airway epithelial cells employ both shared detoxification strategies and cell type-specific adaptive responses to cadmium toxicity.

**Figure 4.**
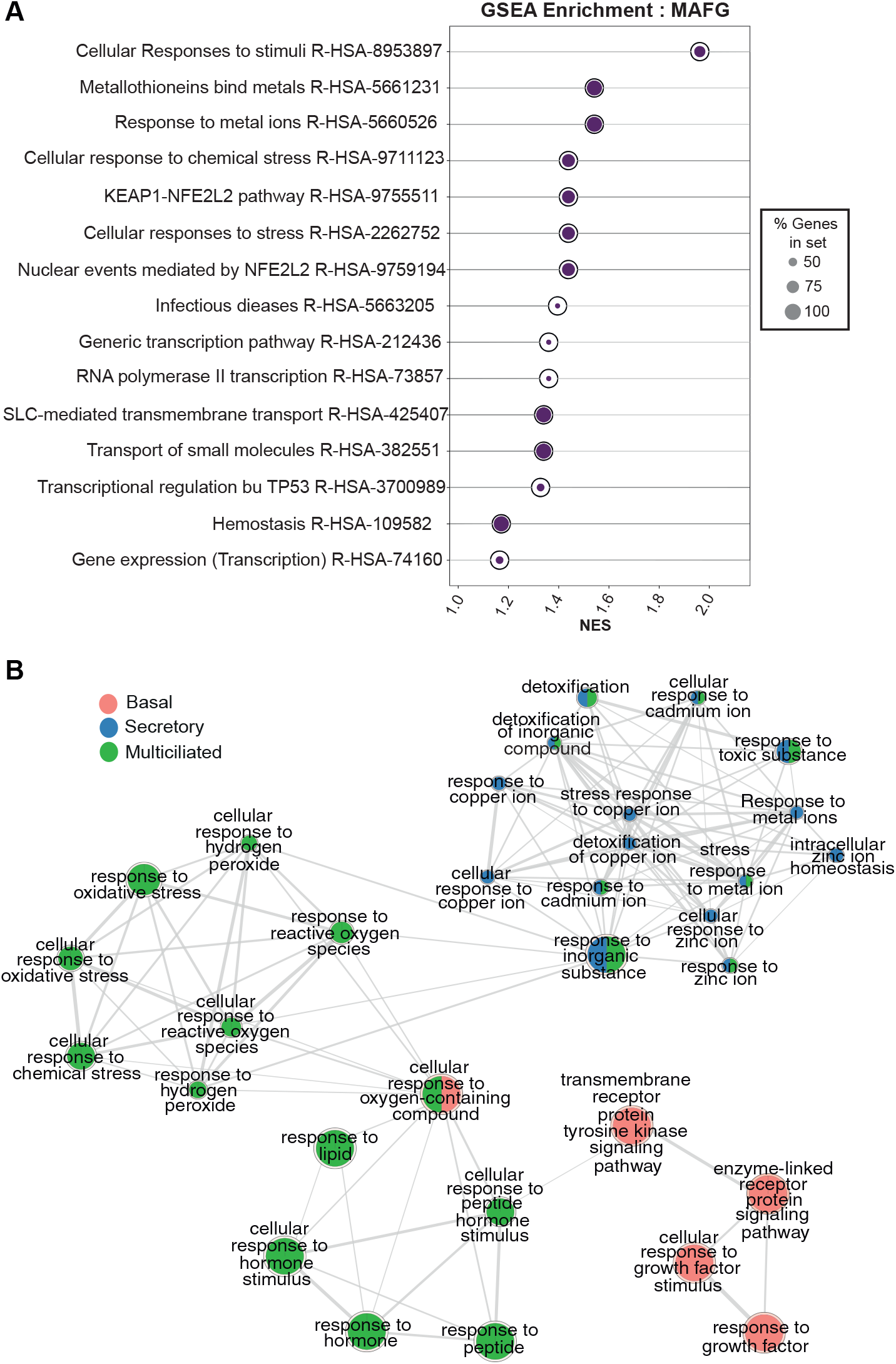
Enrichment Map of Gene Set Network Analysis for Identified Regulons. **(A)** Gene Set Enrichment Analysis (GSEA) of MAFG regulon target genes. Dot plot shows enriched Reactome pathways for genes regulated by the MAFG transcription factor. The x-axis represents the normalized enrichment score (NES), indicating the degree of enrichment. Dot size represents the percentage of genes from each pathway present in the MAFG regulon gene set. Top-enriched pathways include metallothionein-mediated metal binding, cellular responses to metal ions and chemical stress, and the KEAP1-NFE2L2 antioxidant pathway. **(B)** Enrichment map derived from gene set network analysis of target genes from regulons identified in CdCl_2_-treated cells across different cell types. Analysis was performed using g:Profiler, and the network was visualized in Cytoscape (v3.10; see Methods). Nodes represent significantly enriched gene sets (FDR q-value < 0.01) from GO Biological Process and Reactome databases, with node size proportional to gene set size. Edges connect gene sets sharing common genes, forming functionally related pathway clusters. Unconnected or minimally connected nodes (≤1 edge) are hidden for clarity. The map reveals distinct functional modules associated with regulon activity in response to CdCl_2_, including metal ion homeostasis, oxidative stress response, and cellular signaling pathways. The grey-highlighted region emphasizes pathways related to detoxification and stress responses, encompassing cellular defenses against oxidative stress, chemical stress, and cadmium/metal ion exposure. These pathways are predominantly active in secretory and multiciliated cells.

To validate the differential expression of some of the genes that were differentially expressed in a cell-type selective manner by CdCl_2_ exposure, we performed in situ hybridization on formalin-fixed paraffin embedded sections of MB-ALI cultures. Tissue samples were collected after six hours of treatment; however, the epithelial layer exhibited considerable fragility, and sectioning led to tissue disaggregation. The specimens did not retain adequate structural integrity for reliable analysis. This phenomenon is likely to be caused by the disruption of tight junctions by cadmium (Cao et al. 2015). By contrast, a three-hour cadmium treatment yielded ALI culture sections that maintained sufficient structural integrity to allow for histological analysis. These sections were stained with a KRT5 probe (in green) to identify basal cells and with an anti-acetylated tubulin (in white) to visualize cilia on multiciliated cells. scRNA-seq revealed a preferential expression of IL1RN in basal cells upon cadmium exposure, although substantial expression in secretory cells was also detected. In situ hydridization confirmed this observation, with staining associated with KRT5-positive cells as well as some staining detected in the middle layer containing secretory cells (**Fig. 5A**). In cadmium-treated MB-ALI cultures, SYTL5 and WIPF3 were predominantly expressed in secretory (**Fig. 5B**) and multiciliated cells (**Fig. 5C**), respectively, while MT1G expression was confined to these two cell populations. In situ hybridization further confirmed the absence of signal in KRT5-positive cells for all three probes (**Fig. 5B, 5C, 5D**).

**Figure 5.**
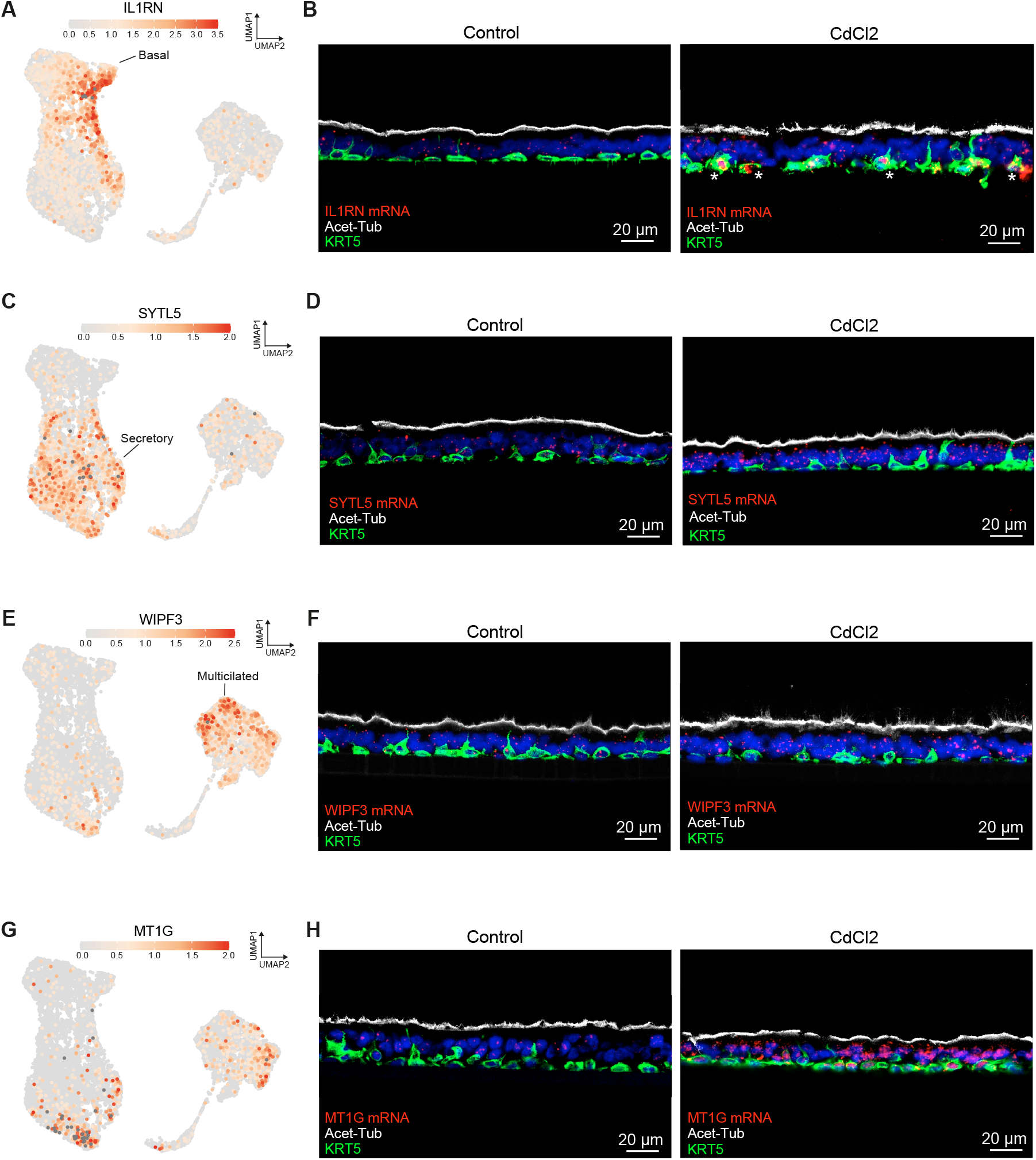
In situ hybridization of cadmium-response genes in MB-ALI cultures. ALI cultures were treated for 3 hours with CdCl_2_ or DMSO. Section of ALI cultures were prepared from paraffin blocks and stained with a KRT5 probe (in green) to identify basal cells and with an anti-acetylated tubulin (in white) to visualize cilia on multiciliated cells. Probes were applied to detect IL1RN (A), SYTL5 (B), WIPF3 (C) and MT1G (D) (all in red). White bar: 20 µm.

## DISCUSSION

We used scRNA-seq to analyze the effects of acute CdCl_2_ exposure on ALI cultures of human bronchial epithelial cells. scRNA-seq data can be analyzed using the pseudobulk method. This approach aggregates single-cell data by summing or averaging counts across cells within defined cell populations, effectively reducing noise and making the data more comparable to bulk RNA-seq. This method is particularly useful for identifying differential gene expression at the population level, as it leverages robust statistical frameworks developed for bulk data. Analysis in pseudobulk of the bulk signatures obtained using differential gene expression analysis between control and CdCl_2_-treated cultures identified a classic response to heavy metals, a modification of the extracellular matrix, a cellular stress response, and the induction of inflammation. These types of responses were expected and have already been described (Forcella et al. 2022; Gasser et al. 2022a; Jomova et al. 2025). The signatures of each cell type make up the bulk signature, but the contribution of each cell type to specific canonical pathways or upstream regulators observed in the bulk signature varies. For example, activation of inflammation was observed in all three cell types (upstream regulator TNF **Fig.2G**). By contrast, the canonical signature related to response to heavy metals was essentially detected in secretory and multiciliated cells (terms Metallothioneins binds metals and metoxalin gadolinium, **Fig 2 G & H**). Cytoskeletal reorganization (IPA term Rho GTPase cycle) was detected in basal cells (**Fig.2G**). Activation of NFE2L2-regulating antioxidant/detoxification enzymes and activation of hypoxia-inducible factor 1-alpha (HIF1α) signaling were only detected in multiciliated cells (**Fig. 2F**). The latter observation is surprising considering that the cells were cultured in normoxic conditions. This HIF1α activation is likely to be the result of the Cd-induced HIF1α stabilization via lysine-63-linked ubiquitination and ER stress (Aschner et al. 2023). Overall, these results highlight the diversity of response to a toxicant of the cell types composing a relatively complex tissue.

An alternative strategy of analysis to the pseudobulk method is SCENIC. SCENIC identifies, for each cell in the dataset, the modulation of regulons, defined as a group of genes that are regulated together by the same transcription factor or regulatory protein. While pseudobulk is simpler and statistically powerful, SCENIC provides mechanistic insights into gene regulation. Together, these approaches can complement each other, offering both population-level and regulatory-level perspectives in scRNA-seq studies. In our case, both approaches confirmed the differential activation of the heavy metal response in secretory and multiciliated cells compared with basal cells. This differential response may be attributed to a differential influx or efflux of Cd into different cell types. A set of genes encoding proteins proven or suspected to be involved in Cd^2+^ entry/efflux into cells has been established (Thévenod 2010). We examined the expression pattern of these genes in our dataset (**Suppl Fig.4**). The genes SLC11A2 and TRPM7 (proven to be involved in Cd^2+^ influx) and ABCC1 (proven to be involved in Cd^2+^ efflux) displayed a homogeneous pattern of expression across the different cell types (**Suppl Fig.4**), suggesting that selective influx or efflux of Cd^2^ was unlikely to be responsible for the differential response of the different cell types.

The enrichment map analysis provides mechanistic insights into cell-type-specific responses to cadmium toxicity in airway epithelium. The clustering of detoxification pathways across multiple cell types may suggest a conserved, coordinated defense response to cadmium exposure, consistent with cadmium’s known ability to disrupt epithelial barrier function through tight junction disruption (Cao et al. 2015) and to induce cellular stress responses (Jomova et al. 2025). The differential pathway enrichment patterns suggest distinct cellular vulnerabilities and adaptive responses. The activation of receptor tyrosine kinase and growth factor signaling pathways specifically in basal cells is a significant point, as these cells serve as progenitor cells maintaining epithelial homeostasis. FOSL1, for which the promoter is inducible by various growth factors and cytokines (Sobolev et al. 2022), shows activation in basal cells that correlates with the observed enrichment of growth factor signaling pathways. This suggests that cadmium exposure may disrupt normal proliferation and differentiation programs in the stem/progenitor compartment, with potential implications for epithelial regeneration and tissue repair. The preferential activation of oxidative stress response pathways in multiciliated cells aligns with their higher susceptibility to redox perturbations (Jain et al. 2024). This finding is consistent with MAFG regulon activation observed in these cells, as MAFG forms heterodimers with Nrf2 to activate antioxidant response element–mediated cytoprotective genes (Hirotsu et al. 2012; Katsuoka et al. 2005). The Nrf2-MAFG axis represents a key regulatory mechanism for antioxidant gene expression, with MAFG itself being transcriptionally upregulated by oxidative stress through an autoregulatory feedback loop (Katsuoka et al. 2005). This suggests an adaptive response to cadmium-induced reactive oxygen species generation in multiciliated cells. The integration of regulon activity with pathway enrichment reveals a hierarchical regulatory network where master transcription factors coordinate cell-type-specific responses to cadmium stress. The FOSL1-mediated growth factor signaling in basal cells and MAFG-dependent antioxidant responses in multiciliated cells represent distinct but complementary mechanisms for maintaining the epithelium under toxic stress conditions.

To validate the differential expression of certain genes in the different cell types, we performed in situ hybridization. Although this method confirmed the conclusions of the scRNA-seq analysis, it is time consuming and lacks the potential for high-throughput validation. Alternative technique, such as FACS analysis on the dissociated ALI culture, could be envisaged but the process is time consuming, prone to inducing cell damage, and does not satisfactorily address the high-throughput issue. In this context, spatial transcriptomics (Moffitt et al. 2022) could not only validate but also complement scRNA-seq, by confirming cell-type-specific responses and revealing microenvironmental interactions. Spatial transcriptomics could also be used in predictive toxicology to detect cell-type-specific expression of genes that may correlate with adverse outcomes.

In conclusion, our data illustrate that scRNA-seq provides higher resolution transcriptional signatures with unexpected features. This information is likely to have implications in the refinement of AOPs and could provide the basis for new generations of tests in predictive toxicology.

## ACKNOWLEDGEMENTS

The authors thank the technical support of the CoBioDA bioinformatics hub, the UCA GenomiX platform and the microscopy facility (part of the « Microscopie Imagerie Cytométrie Azur» GIS IBiSA labelled platform) at IPMC. We acknowledge support from the Centre National de la Recherche Scientifique (CNRS), Institut National de la Santé et de la Recherche Médicale (Inserm), Université Côte d’Azur. This work was supported by a grant from the French government managed by the Agence Nationale de la Recherche under the France 2030 programme, reference ANR-23-IAHU-0007, the Agence Nationale de sécurité sanitaire (AAP PNR EST 0812) and BPI France (grant DOS0177167/00).

## STATEMENTS AND DECLARATIONS

The authors declare that they have no conflict of interest.

## DATA AVAILABILITY

The dataset that supports the findings in this study is openly available (GEO accession number GSE316799).

## Notes

### Competing Interest Statement

The authors have declared no competing interest.

