## Supplementary material for "Single-cell transcriptomics reveals a differential response of human bronchial epithelial cell-types to cadmium chloride": Suppl Fig 1

A

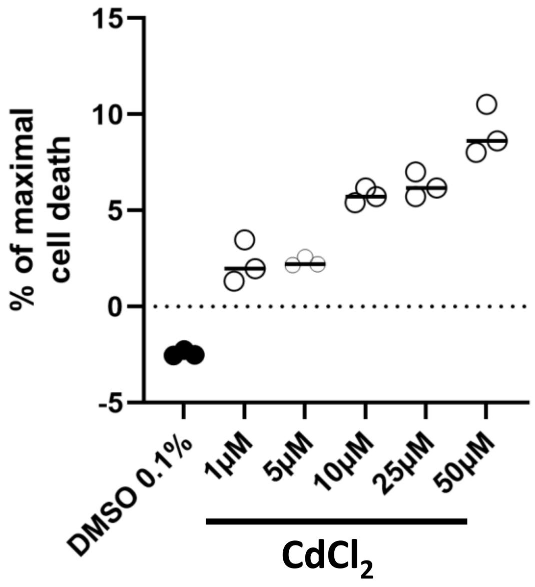


B

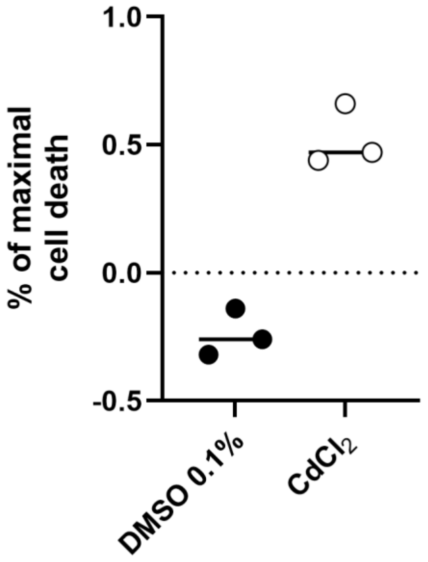


**Supplemental Fig.1: Determination of the toxicity of CdCl2 on pulmonary cells.** A) Beas-2B cells (10000) were seeded in a 96 well plate. The next day, the medium was changed and supplemented with various concentrations of CdCl2. B) ALI cultures were exposed for 24 hours with 10µM CdCl2. In both experiments, medium (200 µL) was collected and the LDH activity was measured. For both dataset, data are expressed as percentage of LDH release induced by incubating cultures with a lysis buffer that induce cell death and maximal LDH release (provided in the CyQUANT kit).
