## Supplementary material for "Single-cell transcriptomics reveals a differential response of human bronchial epithelial cell-types to cadmium chloride": Suppl Fig 2

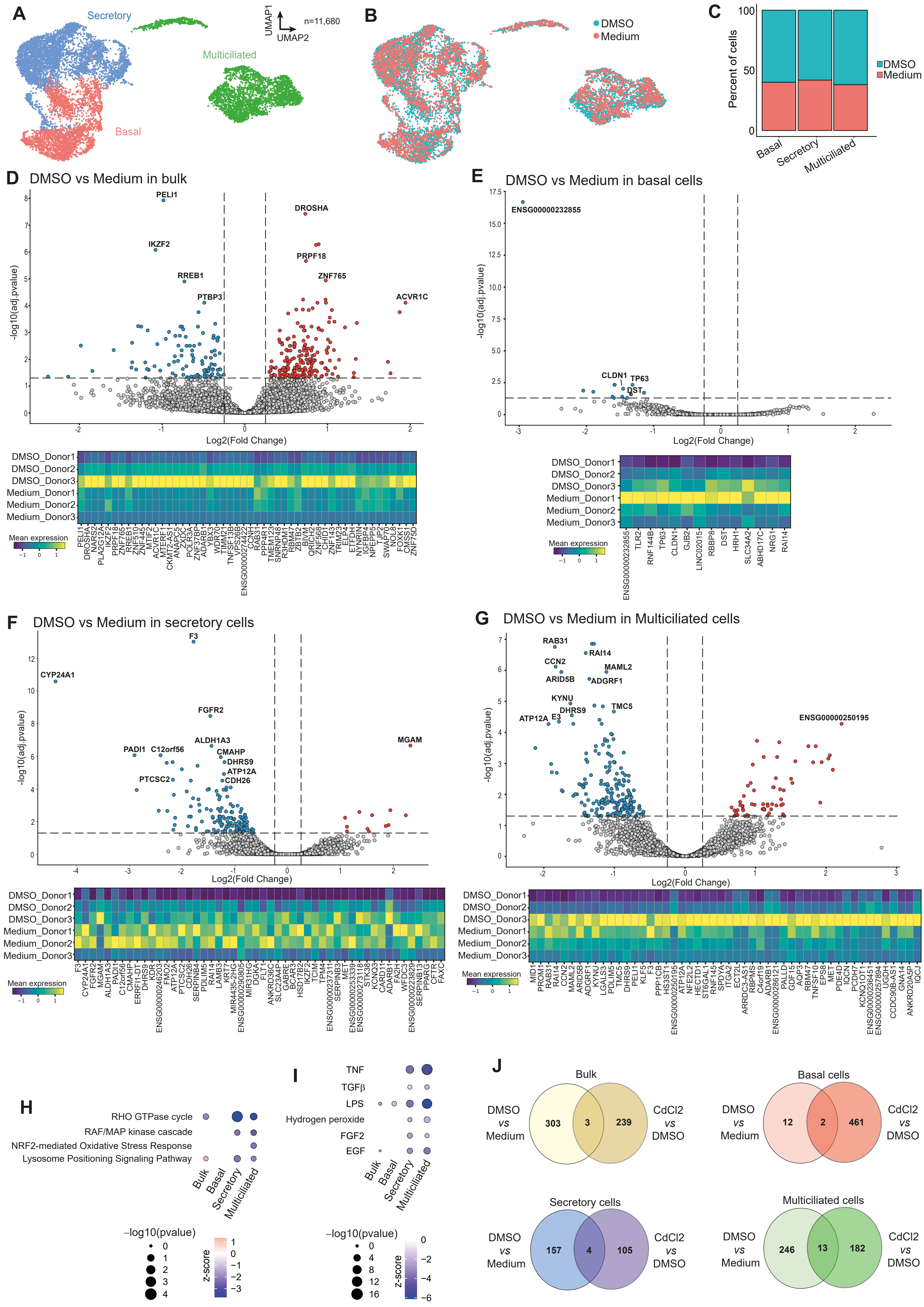

A
Secretory
Multiciliated
Basal
UMAP1
UMAP2
n=11,680
B
DMSO
Medium
C
Percent of cells
100
50
0
Basal
Secretory
Multiciliated
DMSO
Medium
D
DMSO vs Medium in bulk
-log10(adj.pvalue)
Log2(Fold Change)
PELI1
IKZF2
RREB1
PTBP3
DROSHA
PRPF18
ZNF765
ACVR1C
DMSO_Donor1
DMSO_Donor2
DMSO_Donor3
Medium_Donor1
Medium_Donor2
Medium_Donor3
Mean expression
E
DMSO vs Medium in basal cells
-log10(adj.pvalue)
Log2(Fold Change)
ENSG00000232855
CLDN1
TP63
DST
ENSG00000232855
TLR2
RNF144B
TP63
CLDN1
GJB2
LINC02015
RBBP8
DST
HRH1
SLC34A2
ABHD17C
NRG1
RAI14
Mean expression
F
DMSO vs Medium in secretory cells
-log10(adj.pvalue)
Log2(Fold Change)
F3
CYP24A1
FGFR2
ALDH1A3
PADI1
C12orf56
CMAHP
DHRS9
ATP12A
CDH26
PTCSC2
MGAM
Mean expression
G
DMSO vs Medium in Multiciliated cells
-log10(adj.pvalue)
Log2(Fold Change)
RAB31
RAI14
CCN2
MAML2
ARID5B
ADGRF1
KYNU
DHRS9
TMC5
ATP12A
E3
ENSG00000250195
Mean expression
H
RHO GTPase cycle
RAF/MAP kinase cascade
NRF2-mediated Oxidative Stress Response
Lysosome Positioning Signaling Pathway
Bulk
Basal
Secretory
Multiciliated
−log10(pvalue)
z-score
I
TNF
TGFβ
LPS
Hydrogen peroxide
FGF2
EGF
Bulk
Basal
Secretory
Multiciliated
−log10(pvalue)
z-score
J
Bulk
DMSO vs Medium
303
3
239
CdCl2 vs DMSO
Basal cells
DMSO vs Medium
12
2
461
CdCl2 vs DMSO
Secretory cells
DMSO vs Medium
157
4
105
CdCl2 vs DMSO
Multiciliated cells
DMSO vs Medium
246
13
182
CdCl2 vs DMSO

**Supplemental Figure 2. Bulk and cell-type specific transcriptional analysis of 0.1% DMSO exposure in human bronchial epithelial cells.**

(A) UMAP plot presenting the different cell-types in the dataset. (B) UMAP presenting the distribution of cell types in the different conditions. (C) Proportions of the different cell-types in medium or 0.1% DMSO-exposed cultures. Volcano plots of and heatmap per donor of the differentially-expressed genes (0.1% DMSO versus medium) in the bulk (D), basal (E), secretory (F) and multiciliated cells (G). Ingenuity Pathway Analysis (IPA) of the canonical pathways (H) and upstream regulators (I) modulated by 0.1% DMSO exposure in the different cell-types. (J) Venn diagrams of the differentially-expressed genes in 0.1% DMSO versus medium and $CdCl_2$ versus 0.1% DMSO in the bulk and the three cell types.
