## Supplementary figures and images for "Single-cell transcriptomics reveals a differential response of human bronchial epithelial cell-types to cadmium chloride"

### Suppl Fig 3

#### Supplemental Figure

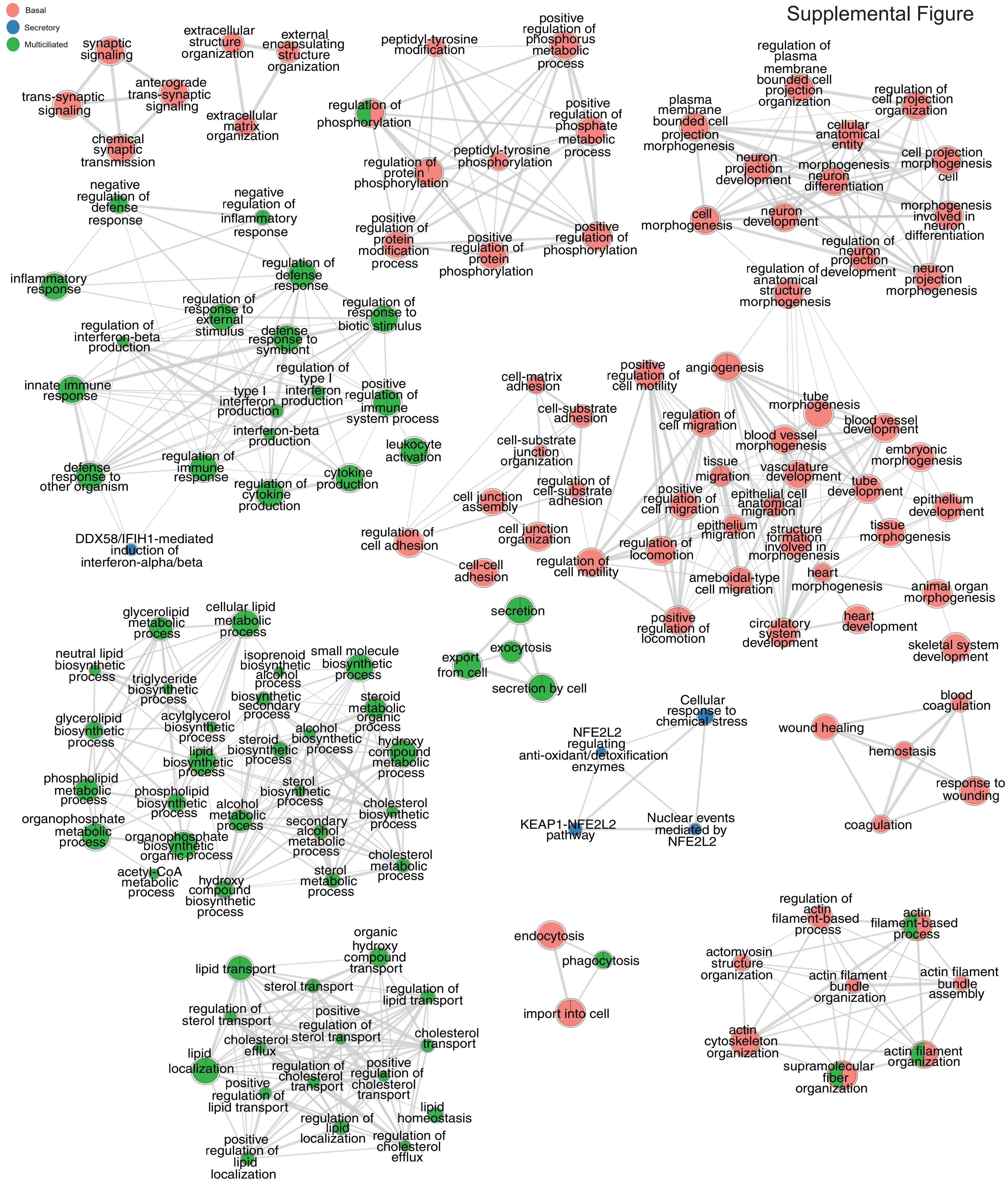

### Suppl Fig 4

##### A Genes involved in the entry of $Cd^{2+}$ into the cells

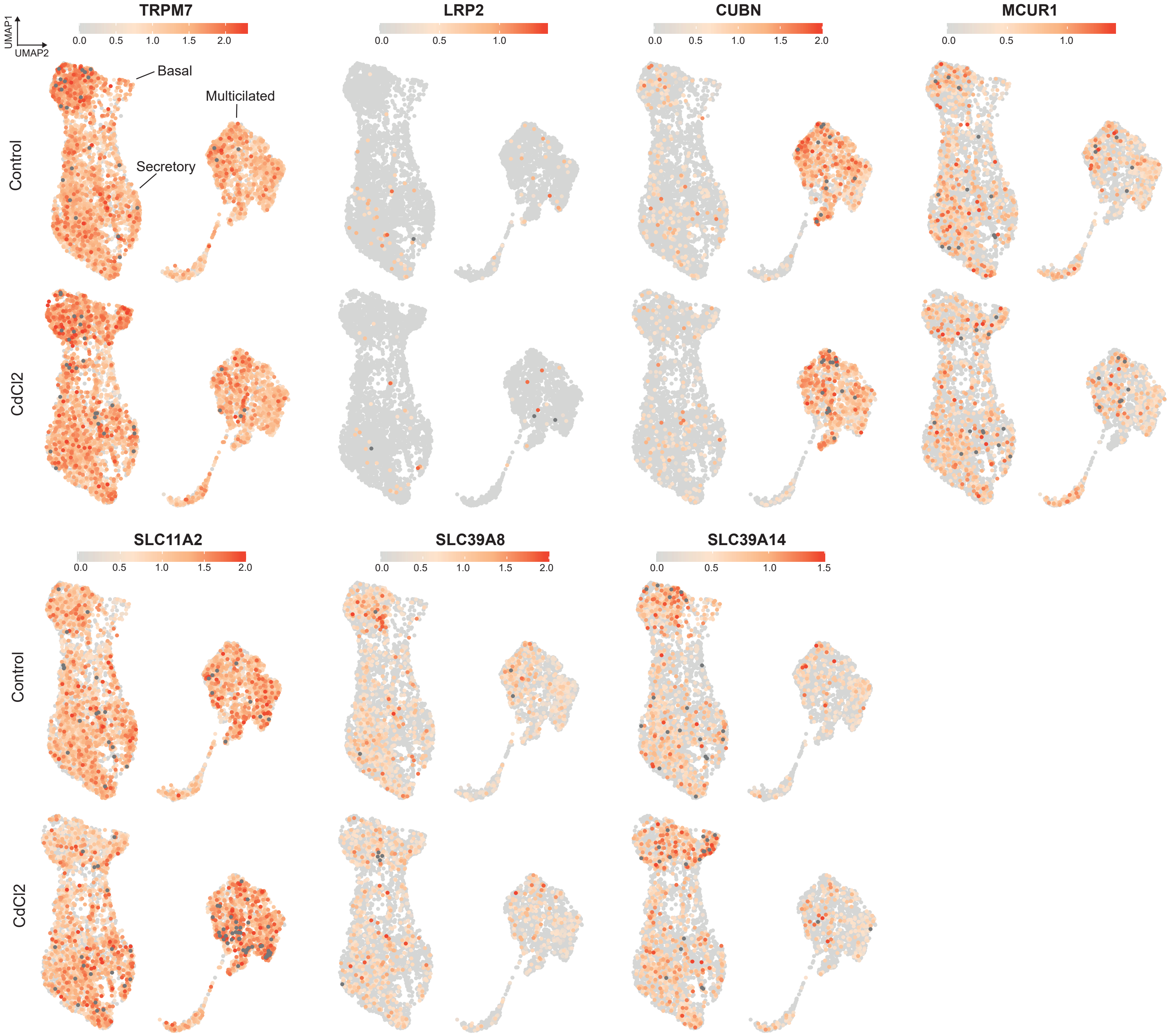


##### B Genes involved in $Cd^{2+}$ efflux from the cells

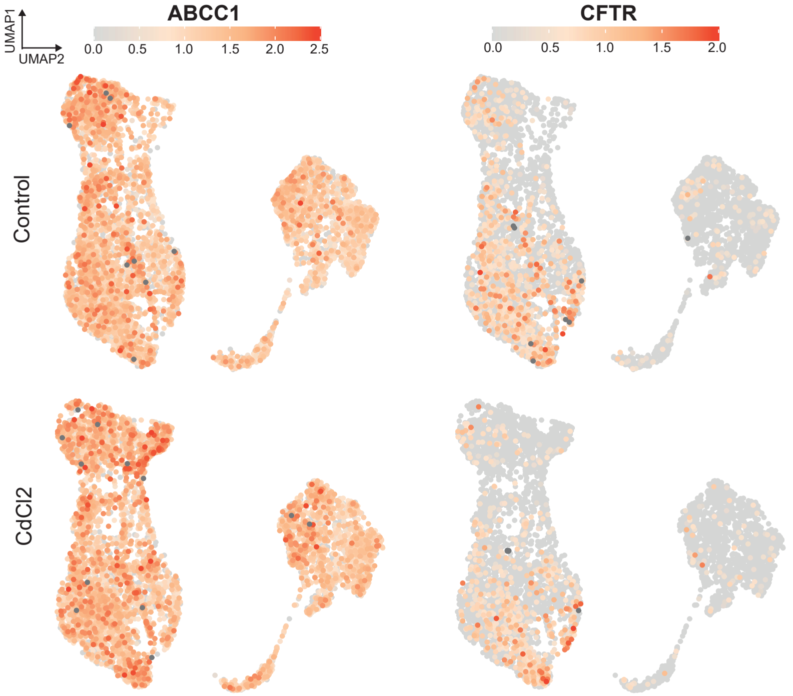
